# Characterization of the early response of *Arabidopsis thaliana* to *Dickeya dadantii* infection using expression profiling

**DOI:** 10.1101/415380

**Authors:** Frédérique Van Gijsegem, Frédérique Bitton, Anne-Laure Laborie, Yvan Kraepiel, Jacques Pédron

**Affiliations:** Sorbonne Université, Institute of Ecology and Environmental sciences-Paris, 4 place Jussieu, F-75 252 Paris, France; INRA, AgroParis Tech, Interactions Plantes-Pathogènes (UMR217), 16 rue Claude Bernard, F-75231 Paris CEDEX 05, France; Unité de Génétique et d’Amélioration des Fruits et Légumes, F-84143 Montfavet, France

**Author notes:** YK and JP contributed equally to this work. Correspondence to: Jacques Pédron, Yvan Kraepiel.

**Keywords:** Arabidopsis thaliana, Dickeya dadantii, Erwinia chrysanthemi, transcriptome analysis, indole glucosinolates, jasmonates, soft rot disease

## Abstract

To draw a global view of plant responses to interactions with the phytopathogenic enterobacterale *Dickeya dadantii*, a causal agent of soft rot diseases on many plant species, we analysed the early *Arabidopsis* responses to *D. dadantii* infection. We performed a genome-wide analysis of the *Arabidopsis thaliana* transcriptome during *D. dadantii* infection and conducted a genetic study of identified responses.

A limited set of genes related to plant defence or interactions with the environment were induced at an early stage of infection, with an over-representation of genes involved in both the metabolism of indole glucosinolates (IGs) and the jasmonate (JA) defence pathway. Bacterial type I and type II secretion systems are required to trigger the induction of IG and JA-related genes while the type III secretion system appears to partially inhibit these defence pathways. Using *Arabidopsis* mutants impaired in JA biosynthesis or perception, we showed that induction of some IG metabolism genes was COI1-dependent but, surprisingly, JA-independent. Moreover, characterisation of *D. dadantii* disease progression in *Arabidopsis* mutants impaired in JA or IG pathways showed that JA triggers an efficient plant defence response that does not involve IGs.

The induction of the IG pathway by bacterial pathogens has been reported several times *in vitro*. This study shows for the first time, that this induction does indeed occur *in planta*, but also that this line of defence is ineffective against *D. dadantii* infection, in contrast to its role to counteract herbivorous or fungal pathogen attacks.

## Introduction

Plants are continuously exposed to pathogen attack and must rapidly recognize the presence of pathogens to initiate a timely, accurate and effective immune response. This immune response includes multiple lines of defence such as production of reactive oxygen species (ROS), plant cell wall reinforcement, synthesis of antimicrobial metabolites and pathogenesis-related (PR) proteins [1]. Many of these defence mechanisms are orchestrated by a complex interplay between signalling molecules of which salicylic acid (SA), jasmonate (JA) and ethylene (ET) are particularly important [2]. These responses are often accompanied by a massive transcriptional reprogramming that favours defence responses over routine cellular requirements. Thus, in *Arabidopsis*, up to 25% of all genes respond to various pathogen infections by altering their transcript levels [3, 4]. Using genome-wide technologies, it is now possible to address the molecular basis of the complex plant response to pathogen attack.

*Dickeya dadantii* (formerly *Erwinia chrysanthemi*), a broad host-range phytopathogenic enterobacterale, is a causal agent of soft rot disease on many crops and ornamentals and on the model plant *Arabidopsis thaliana* [5, 6]. *D. dadantii* virulence relies mainly on the production and secretion of plant cell wall degrading enzymes into the extracellular spaces of infected tissues [7–9]. These include pectinases and a cellulase secreted by a type II Out secretion system (T2SS) as well as proteases secreted by a type I Prt secretion system (T1SS) [10, 11]. Other plant factors influence efficient colonization, such as bacterial iron uptake systems, Hrp type III secretion system (T3SS), resistance to chemical and environmental stresses encountered within plant tissues, bacterial envelope constituents or motility [8, 9, 12, 13, 14, 15, 16]. Often, after invading its host plant, *D. dadantii* resides latently in the plant intercellular spaces without provoking any symptoms, exhibiting a biotrophic lifestyle characterized by the repression of genes encoding virulence factors by several bacterial global regulators [17, 18, 19]. As for hemibiotrophic and most bacterial pathogens [20], disease only occurs when bacteria shift to a necrotrophic behaviour under environmental conditions - such as high temperatures, high humidity - favourable for massive bacterial multiplication and production of virulence factors [15, 21].

Plant defence responses provoked by soft rot enterobacterales have been mainly studied using *Pectobacterium carotovorum subsp. carotovorum* and *D. dadantii* on different host plants. Identified plant defence responses to *D. dadantii* infection are multiple, but nevertheless non decisive - no monogenic resistance to *D. dadantii* has been characterized - and perturbations of individual plant responses have a weak impact on symptom severity. These responses include competition for iron availability in *Arabidopsis* [6, 22, 23], activation of defence-related ELI genes in parsley [24] and triggering of early extracellular ROS production associated with plant cell wall reinforcement [25, 26, 27, 28]. Both ROS production and cell wall reinforcement increased in ABA deficient mutants during *D. dadantii* infection [26, 29]. Modulation of plant cell death (PCD) also participates to the *D. dadantii*/*Arabidopsis* interaction, since exacerbated necrosis during infection limits disease progression [27]. Similarly, a pathogen-triggered inhibition of PCD was also identified during the *P. carotovorum subsp. carotovorum*/*Arabidopsis* interaction [30]. In both *P. carotovorum subsp. carotovorum* and *D. dadantii* species, bacterial infection provoked the activation of plant defence marker genes associated to SA and JA/ET signalling pathways [25, 31, 32, 33, 34]. The JA deficient *jar1* mutant was shown to be more susceptible to *D. dadantii* infection, whereas the SA deficient *sid2* mutant was as susceptible as the wild type [25], indicating that JA but not SA-dependent defence is partially efficient against *D. dadantii* invasion. Nevertheless, Antunez-Lamas et al. [34] reported that JA plays a “dual role”, as it also promotes bacterial entry through wounded tissues.

These data show that the *D. dadantii*/*Arabidopsis* interaction is a complex process involving multiple plant responses. Among these defence responses, some are partially efficient (such as ROS accumulation or the JA pathway), some are not (SA pathway), and conversely, some promote disease progression (such as inhibition of PCD, ABA pathway). As none of these plant responses is decisive in the disease issue, we hypothesised that other not already examined plant responses could occur during the interaction. In order to identify such new plant factors involved in this interaction, regardless the role - whether positive or negative - of these factors, we adopted a without *a priori* strategy based on the genome scale transcriptome analysis of *Arabidopsis* during the early stage of *D. dadantii* infection. This study showed increased expression levels of three sets of genes during the early stage of infection. These included genes involved in the biosynthesis of tryptophan, genes involved in the metabolism of indole glucosinolates (IGs) tryptophan-derived secondary metabolites and genes involved in the JA biosynthesis and signalling pathway. By using bacterial or plant mutants, we further analysed the relationship between JA and IG pathways, the nature of the bacterial triggering factors and the efficiency of these responses on disease progression.

## Material and methods

### Biological material and growth conditions

*D. dadantii* wild type and mutant strains used in this study are listed in Supplemental Table S1. The wild type strain 3937 (our collection) was isolated from *Saintpaulia ionantha* (African violet). Bacterial strains were grown at 30°C in Luria-Bertani medium. Liquid cultures were grown in a shaking incubator (180 rpm). For plant inoculations, the bacterial inoculum was prepared as follows: an aliquot of the −80°C glycerol stock culture was streaked on solidified LB medium (1.5% Difco agar) and grown for 48 h at 30°C. Then, a single colony was used to inoculate a liquid culture. After 8 h of growth at 30°C, 100 μl of the culture was plated on LB agar medium and incubated overnight. The bacteria were then suspended in inoculation buffer (50 mM KPO_4_, pH 7) and inoculum density was adjusted to the value mentioned in each experiment.

*Arabidopsis thaliana* lines used in this study are listed in Supplemental Table S1. *Arabidopsis* seeds were sown in non-sterile soil, synchronized by incubation for two days at 4°C and allowed to germinate and grow for three weeks. They were then potted individually (pathogenicity assays) or in threes (RNA isolation) into separate pots and incubated for a further three weeks before inoculation. Plants were grown under short day condition (8h day/16h night) at 24°C/19°C.

### Plant inoculation

Six-week-old plants were heavily watered and covered with a transparent lid 16 h before inoculation. The cover was kept in place throughout the assay to maintain high-moisture conditions.

For plant RNA isolation, plants were infected by rapid immersion in a bacterial suspension (5. 10^7^ cfu/mL) prepared in inoculation buffer containing 0.01% (vol/vol) Silwet L-77 surfactant (van Meeuwen Chemicals BV, Weesp, The Netherlands). Aerial plant tissues were then collected either immediately after immersion (0 hpi) or 6, 12 and 24 hours post inoculation (hpi) and ground in liquid nitrogen to a fine powder.

For pathogenicity assays, plants were infected by wounding one leaf per plant with a needle and depositing a 5μl droplet of a 10^4^ cfu mL^-1^ bacterial suspension on the wound (50 bacteria per droplet). Wounding was necessary to obtain sufficient symptoms with respect to the low density inoculum used and to synchronize symptom outbreaks. This second method of plant inoculation was chosen to allow the scoring of individual symptom progression from 1 to 5 days. Assays were carried out at least in duplicates. Statistical significance analysis of the results was performed using Fisher’s exact test, with a 0.05 two sided p-value threshold.

### RNA isolation for CATMA array hybridization and for quantitative Real-Time PCR

Total RNAs were purified by extraction in a guanidinium isothiocyanate extraction buffer and recovery by centrifugation on a caesium chloride cushion as described in [17].

### Transcriptome CATMA array hybridization

Transcriptome profiling was carried out at the URGV - Plant Genomics Research (Evry, France) using Arabidopsis CATMA arrays containing 24,576 gene-specific tags (CATMAv2 repertoire, [35, 36]). Arrays were printed and hybridized as described by Lurin *et al.* [37]. Each experiment was carried out with two independent biological repeats to take into account biological variation. Each comparison included a technical repeat with fluorochrome reversal (dye swap). RNA integrity was verified using a Bioanalyzer (Agilent, www.agilent.com). For each sample, cRNAs were synthesized from 2 μg of total RNA using the Message Amp aRNA kit (Ambion, www.ambion.com), and 5 μg of cRNA was reverse transcribed using 300 units of SuperScript II (Invitrogen) and Cy3-dUTP or Cy5-dUTP (NEN) for each slide. Labelled samples were combined, purified and concentrated using YM30 Microcon columns (Millipore). The hybridization of labelled samples to the slides and the scanning of the slides were performed as described by Lurin *et al.* [37].

### CATMA array data analysis

Transcriptome statistical analysis was based on dye swaps (i.e. two arrays each containing 24,576 GSTs and 384 controls). For each array, the raw data consisted of the log2 median feature pixel intensity without background subtraction at 635 nm (red) and 532 nm (green). Data analysis was performed as described by Gagnot *et al*. [38]. In the following description, log ratio refers to the differential expression between two conditions: either log2 red/green or log2 green/red according to the experimental design. Array-by-array normalization was performed to remove systematic biases: spots with poor features were excluded. To correct for dye bias, the signal was normalized globally according to intensity using the LOESS procedure. For each block, the log ratio median calculated over the values for the entire block was subtracted from each individual log ratio value to correct for print tip effects in each metablock. To determine differentially expressed genes, the log ratios were analysed using a paired t-test, assuming that the variance of the log ratios is the same for all genes. Spots with extreme variance (high or low) were excluded. The raw P values were adjusted by the Bonferroni method (type I error equal to 5%) to control for family-wise error rate and to drastically limit false-positives in a multiple comparison context [39].

### Quantitative real-time reverse transcription-polymerase chain reaction (qRT-PCR)

RNA samples were treated with RNAse-free DNAse I (Invitrogen) to remove any DNA contamination. First-strand cDNAs were then synthesized from 1-3 μg of total RNA using oligo(dT20) primer and M-MLV reverse transcriptase (Invitrogen), following the manufacturer’s instructions. For quantitative Real-Time PCR analysis, cDNAs were amplified using Maxima^®^ SYBR Green/ROX qPCR Master Mix (Fermentas) according to the manufacturer’s license in an Applied Biosystems 7300 Real Time PCR System using the following conditions: 10 min at 95°C followed by 40 amplification cycles each consisting of 15 s at 95°C and 60 s at 60°C. Results were analysed with the Applied Biosystems Sequence Detection Software v1.3.1.

All primers are listed in supplemental table S1. The *ß-6 TUBULIN* (*TUB6*) gene was used as an internal control and behaved similarly all through the kinetics. This gene was thus used to normalize the expression data for each gene of interest. The comparative quantitation method (ΔΔCt) was used to contrast the different treatments [40]. Ct values quantify the number of PCR cycles necessary to amplify a template to a chosen threshold concentration, ΔCt values quantify the difference in Ct values between a test and a control gene for a given sample, and ΔΔCt values are used for the comparison between two samples. ΔΔCt values were transformed to absolute values with 2^-ΔΔCt^ for obtaining relative fold changes. Relative fold changes for each gene were normalised to 0 hpi control (set to one). All assays were run in duplicates (biological replication) to control for overall variability.

## Results

### *D. dadantii* induces the expression of a limited set of genes during the early phase of *Arabidopsis* infection

To investigate *Arabidopsis* responses to *D. dadantii*, a genome-scale expression profiling was performed using CATMA arrays covering 24, 576 nuclear genes [35, 36]. We focused the study on genes regulated during the early phase of infection. For this purpose, plants were infected by immersion of aerial parts in a bacterial suspension and changes in gene expression were investigated 12 and 24 hours post-inoculation (hpi). Since with this method of inoculation no broad maceration symptoms were observed before 30 hpi, these time points should reflect early plant responses rather than indirect consequences of tissue maceration and plant cell death.

The microarray analysis revealed that a limited set of genes displayed significant differential expression (Bonferroni *P* value, threshold of 5%) during infection. 274 genes (i.e. around 1% of the total protein encoding genes), corresponding to 263 up-regulated and 11 down-regulated genes, displayed significant differential expression 12 or 24 hpi as compared to the mock-inoculated controls (Fig 1A, Supplemental Table S2). Of these genes, 61 were up and 1 was down-regulated at both 12 and 24 hpi.

**Fig 1.**
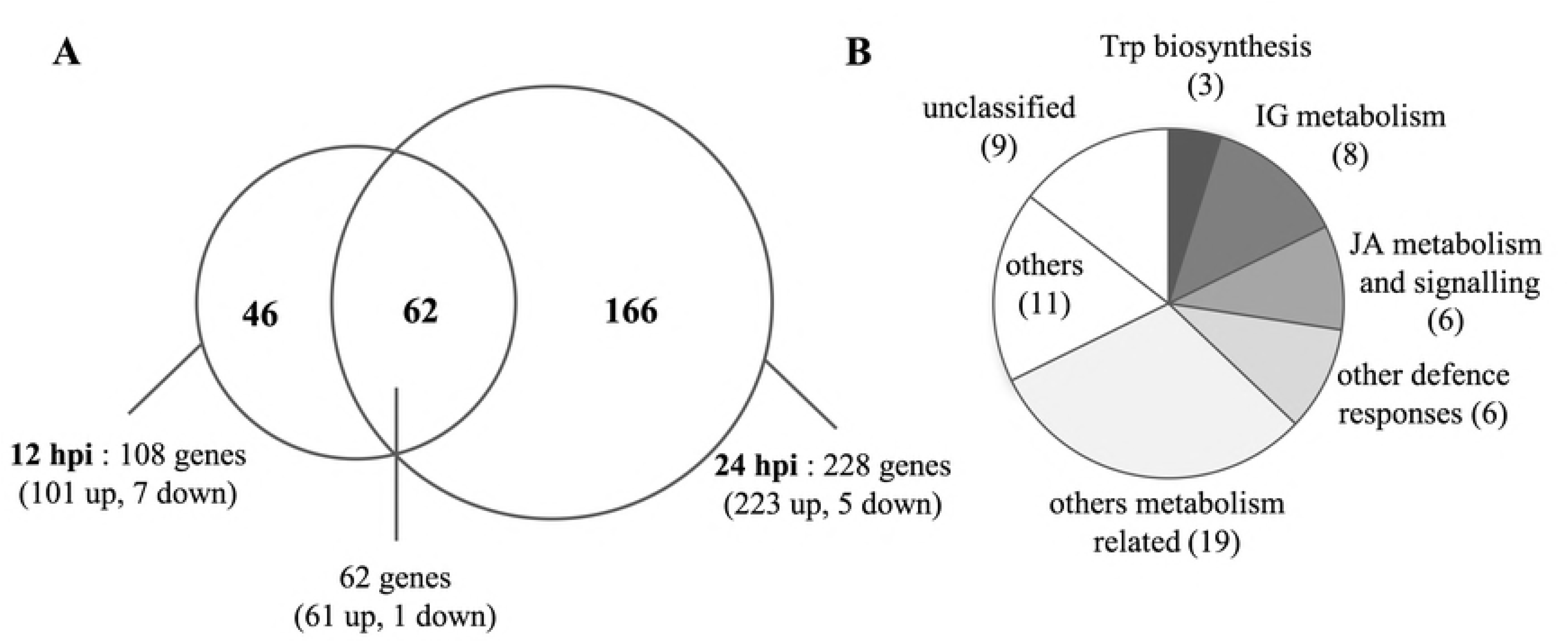
Global analysis of the *Arabidopsis* transcriptome in response to *D. dadantii* infection. A: Venn-diagram of up and down-regulated genes 12 and 24 hours post-inoculation (hpi). B: functional classification of the 61 genes up-regulated both 12 and 24 hpi.

We analysed the putative functions of these genes by using the Protein Sequence database of the Munich Information Centre [41]. 64% of the differentially expressed genes are annotated as encoding proteins of known or predicted functions, distributed among 3 main categories (Table 1, Fig 1B). Not surprisingly, nearly one third of the modulated genes are associated with the “interaction with the environment, cell rescue and defence” category. Another third corresponds to the “metabolism-related” category, with an over-representation of amino acid and secondary metabolisms. The third main category (15%) is related to the “cellular transport” category (electron transport, drug/toxin transport). Down-regulated genes do not clustered in any special functional category.

**Table 1.** Functional classification of *Arabidopsis* genes regulated in response to *D. dadantii* infection. The sum of percentages exceeds 100% as a single same gene may belong to several functional categories.

| Functional category | number of genes | % of total genes |
| --- | --- | --- |
| **Metabolism** | **99** | **36 %** |
| amino acid metabolism | 19 | 7 % |
| C-compound and carbohydrate metabolism | 31 | 11 % |
| secondary metabolism | 20 | 7 % |
| **Cellular transport, transport facilities and transport routes** | **42** | **15 %** |
| **Interaction with the environment, cell rescue, defence and virulence** | **79** | **29 %** |
| cellular sensing and response to external stimulus | 54 | 20 % |
| stress response | 38 | 14 % |
| other plant signalling molecules response (jasmonic acid, | 18 | 7 % |
| salicylic acid etc.) |  |  |
| response to wounding | 17 | 6 % |
| disease, virulence and defence | 15 | 5 % |
| detoxification | 11 | 4 % |
| ethylene response | 5 | 2 % |
| **Unclassified** | **99** | **36%** |

### Induction of tryptophan-related, indole glucosinolate-related and jasmonate-related genes composes an early response to *D. dadantii* infection

Among the 62 genes reproducibly up-regulated both 12 and 24 hpi, more than one half (34 of 62 genes) are classified as metabolism-related, with an over-representation of indole glucosinolate (IG) metabolism genes (8 genes) and of jasmonate (JA, 3 genes) and tryptophan (Trp, 3 genes) metabolism genes. Other metabolism-related genes are scattered across various metabolic processes (amino acids metabolism, cell wall modification, lipids metabolism). Of the entire set of 274 genes modulated either at 12 or 24 hpi, IG, JA and Trp-related genes (metabolism and signalling) represent more than 10% of the modulated genes, with respectively 13, 10 and 7 genes (Table 2), that cover the entire IG metabolic pathway, from the synthesis of the key precursor Trp to almost every biosynthesis and degradation steps (Fig 2). Notably, *D. dadantii* infection also modulates the expression of the gene encoding the MYB51 transcription factor involved in IG metabolic regulation [42]. Up-regulated JA biosynthesis genes are involved in the early steps of JA biosynthesis, with three *LOX* genes (conversion of α-linolenate) and three genes involved in oxo-phytodienoate (OPDA) conversion to OPC8-enoyl-CoA (Table 2). In addition to these biosynthesis genes, four of the modulated genes are involved in JA signalling including two *JAZ* repressors and the *ERF2* transcription factor involved in the perception of JA [43].

**Table 2.**
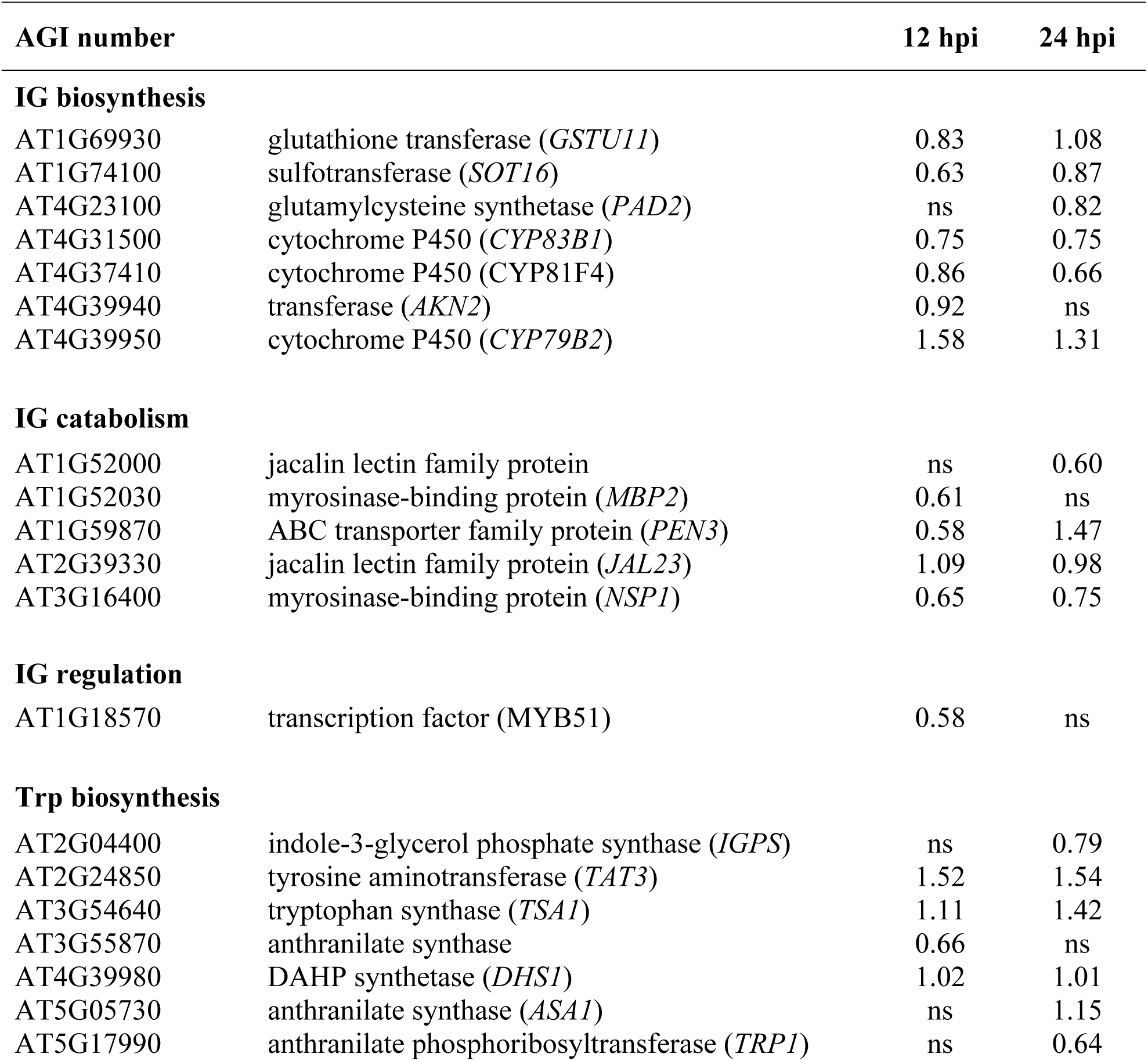
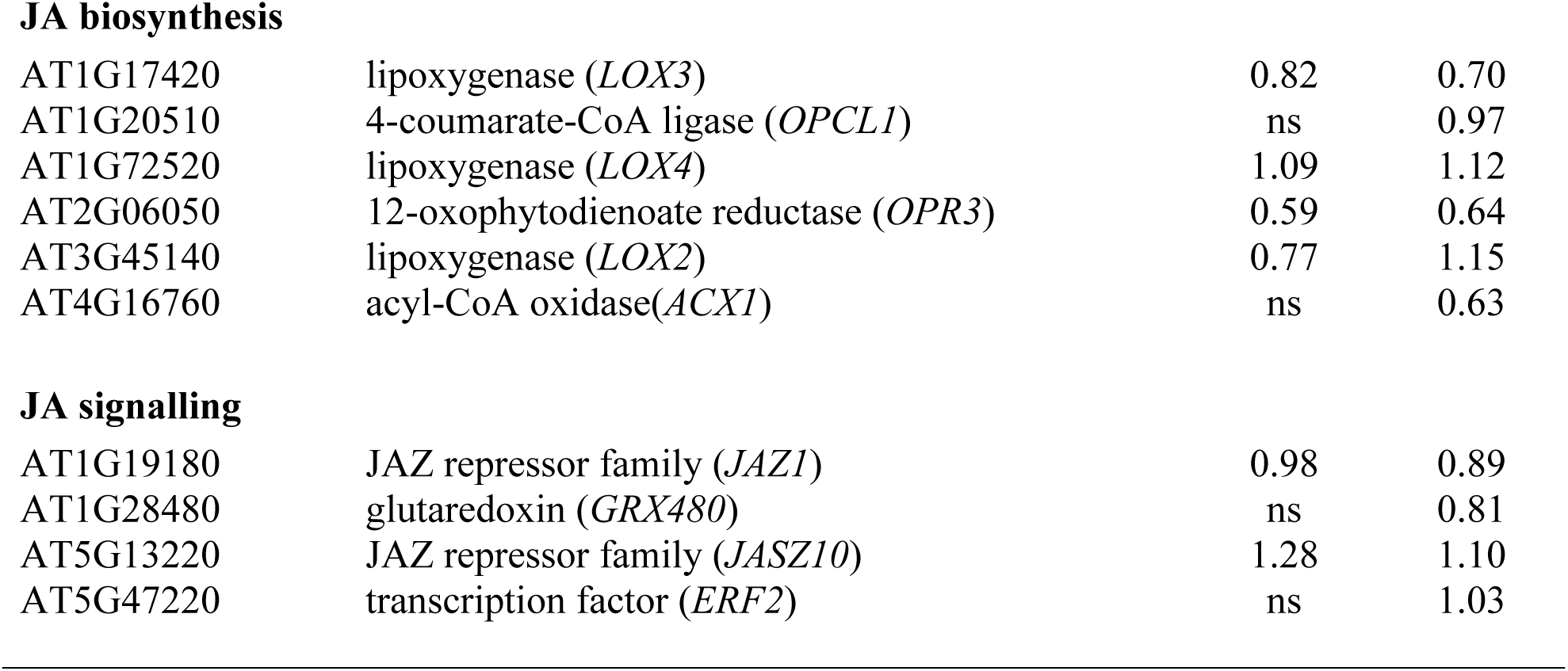
Tryptophan-(Trp), indole glucosinolate-(IG) and jasmonate-(JA) related genes whose expression levels are modulated in response to *D. dadantii* infection. Values are log2 microarray signal ratios between infected and buffer-treated control plants. Positive values correspond to genes up-regulated in response to *D. dadantii* (log2 ratio > 0.5, Bonferroni *P* value < 0.05). ns indicates that gene expression was not significantly changed. hpi: hours post-inoculation.

**Fig 2.**
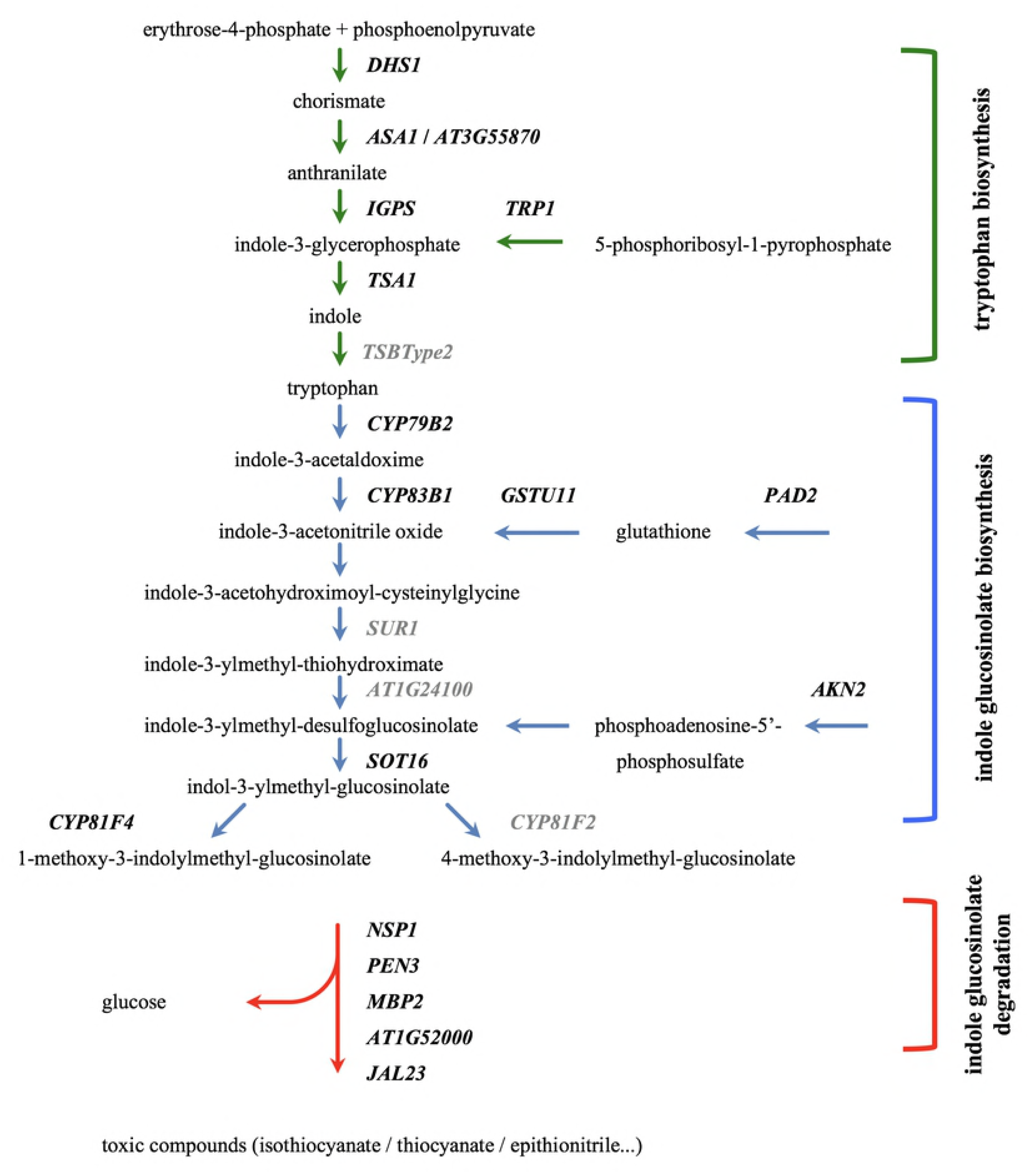
Tryptophan (Trp) and indole glucosinolate (IG) metabolic pathways. All Trp and IG metabolism-related genes in black are up-regulated in response to *D. dadantii* inoculation. Genes in grey were absent from the CATMAv2 array repertoire. Tryptophan is the key precursor of the indole glucosinolates biosynthesis. The last step of the IG biosynthesis catalysed by the *CYP81F4* and *CYP81F2* genes is a diversification step of the final products. Indole glucosinolates are then compartmented. Upon tissue disruption by chewing insects or macergenic pathogens, the action of hydrolases leads to the production of various highly toxic and instable breakdown products (simple nitrile, epithionitrile, thyocyanate) involved in pathogen resistance.

The transcriptomic results were confirmed by qRT-PCR for a subset of Trp, IG and JA-related genes (Fig 3). All gene transcripts progressively accumulated from 12 hpi onwards, with a relative fold change ranging from 1.9 to more than 80 24 hpi. In contrast to the response to the infection by *D. dadantii*, buffer inoculation did not provoke any significant and continued increase of transcripts accumulation. The distribution of experimental values shows that, if a variability of relative fold changes among biological repeats existed, the induction profiles were always similar. This variability was encountered in all the quantitative gene expression analyses we performed.

**Fig 3.**
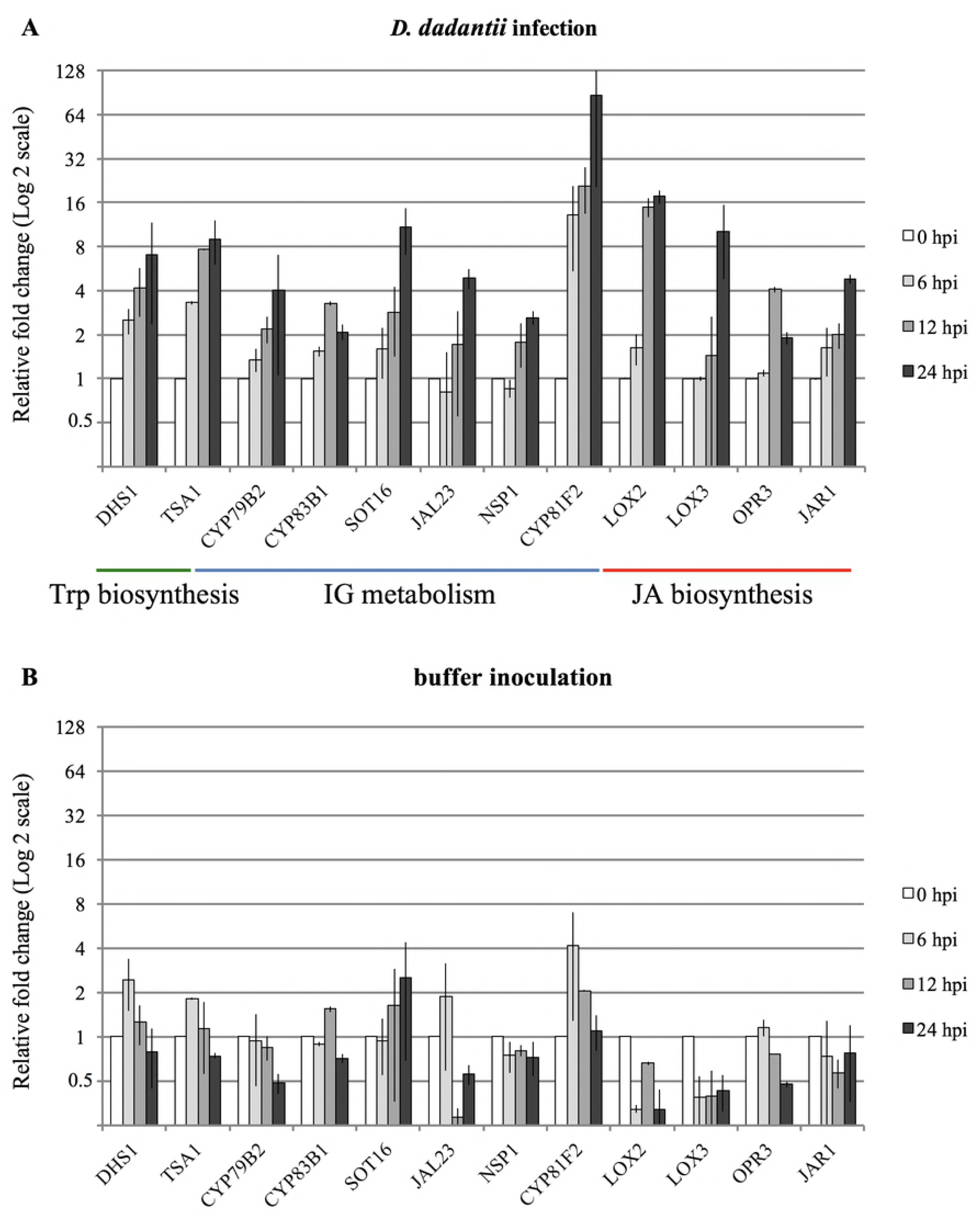
Transcriptome validation by qRT-PCR. Transcript accumulation of representative Trp, IG and JA-related genes after *D. dadantii* (A) or buffer (B) inoculation was analysed by qRT-PCR. Fold changes were normalised to 0 hpi for each gene (set to one). Error bars show the experimental values of the two biological repeats.

### JA mutants have contrasting effects on the IG biosynthetic pathway

Coordinated accumulation of both JA and IG-related gene transcripts during infection prompted us to analyse whether JA plays a role in the control of IG metabolism. This was investigated by analysing the expression of IG biosynthesis genes in JA mutants. The expression of two IG biosynthesis genes - *CYP79B2* (top of the biosynthetic pathway) and *SOT16* (final step of the biosynthetic pathway) - was analysed in the JA insensitive *coi1* mutant [44] and in the JA biosynthesis mutants *jar1* mutant that is impaired in the biosynthesis of the biologically active JA-Ile conjugate [45, 46].

Activation of the two IG biosynthesis genes was not significantly different in the JA biosynthesis mutants *jar1* compared to the col-0 wild type plant 24 hpi (Fig 4). By contrast, in the JA insensitive *coi1* mutant no significant induction of the *SOT16* gene was observed 24 hpi, while the induction of the *CYP79B2* gene was not altered (Fig 4). These results reveal the diversity of the interplay of the JA pathway on IG-related gene regulations: the first step of the biosynthetic pathway seems to be independent of JA biosynthesis and signalling, while the final step of the biosynthetic pathway catalysed by the SOT16 sulfotransferase are regulated by the JA pathway. Interestingly, this regulation involves the JA-Ile receptor *coi1,* rather than JA biosynthesis itself.

**Fig 4.**
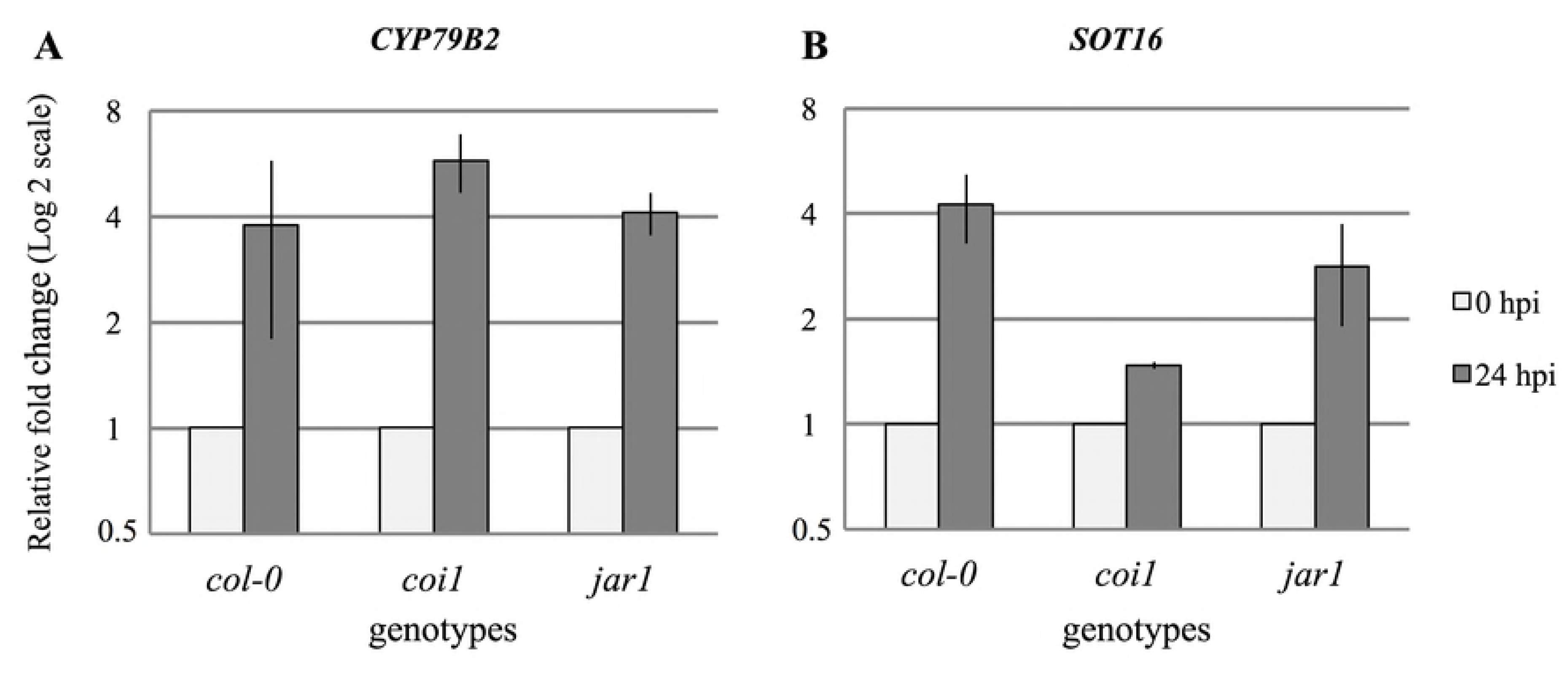
Involvement of the JA pathway on IG biosynthesis gene expression. Transcript accumulation of two representative IG biosynthesis genes *CYP79B2* (A) and *SOT16* (B) was analysed by qRT-PCR 24 hours after *D. dadantii* inoculation on Col-0 wild type *Arabidopsis* and mutants impaired in JA biosynthesis (*jar1*) or impaired in JA sensitivity (*coi1*). Fold changes were normalised to 0 hpi for each *Arabidopsis* genotype (set to one). Error bars show the experimental values of the two biological repeats.

### Type I (PrtE) and type II (OutC) bacterial protein secretion systems are necessary to induce JA and IG-related gene expression during *D. dadantii* infection

Bacterial secretion systems are known to be critical in many plant/bacteria interactions. To test the involvement of these systems on plant transcript accumulation after *D. dadantii* infection, we analysed the requirement for individual bacterial secretion systems in the induction of IG- and JA-related genes. For this analysis, we extended the infection kinetics up to 30 hpi when the variability of inductions was reduced compared to the earlier time points. This allows us to outline an accurate picture of gene induction according to bacterial genotypes.

We quantified by qRT-PCR transcript accumulation levels of two IG-related genes (Fig 5A and B) and two JA-related genes (Figs 5C and D) after buffer inoculation, inoculation with the wild type *D. dadantii* strain (3937) and with mutants altered in the three protein secretion systems known to be involved in virulence (*prtE*, type I mutant, *outC*, type II mutant and *hrcC*, type III mutant).

**Fig 5.**
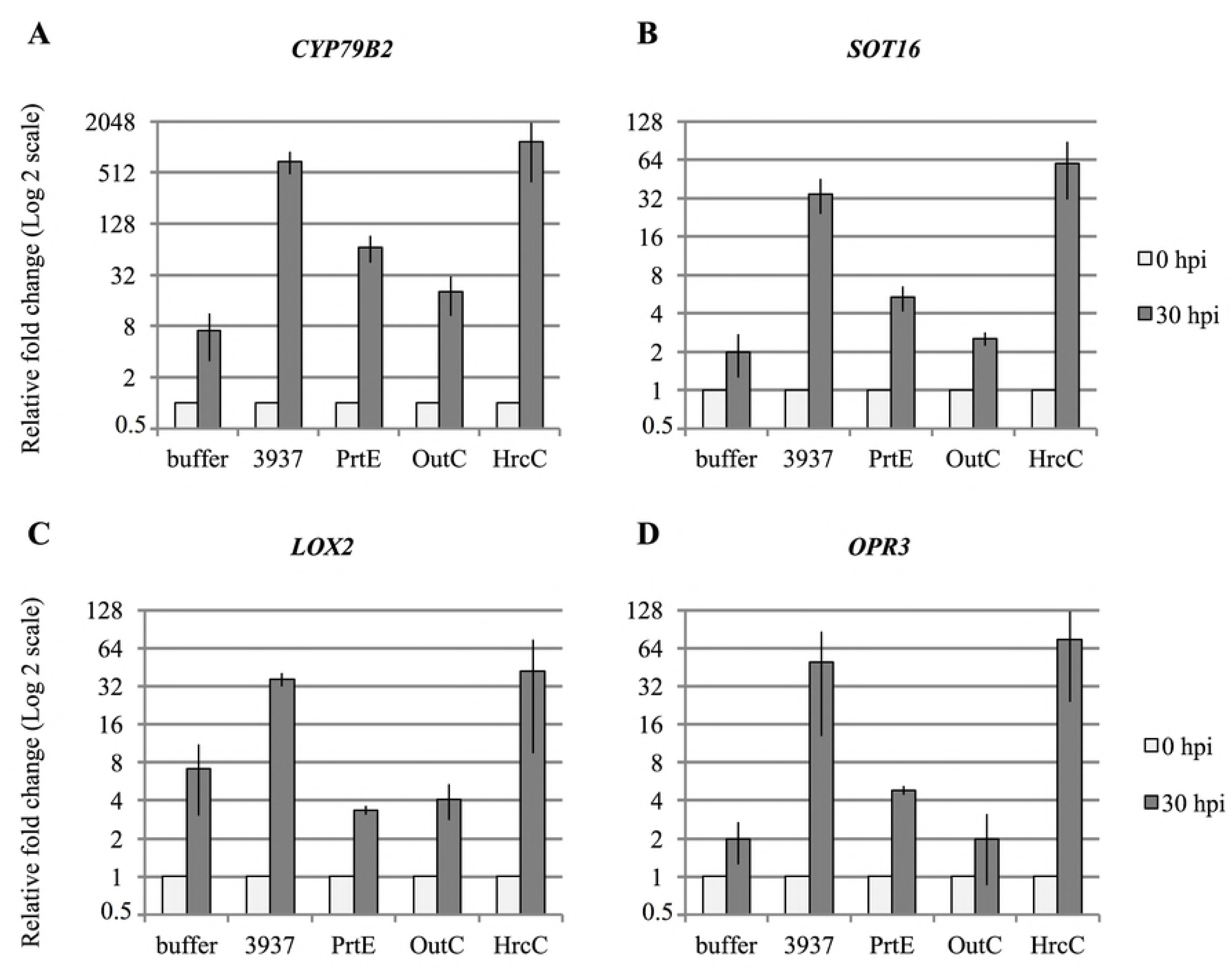
Analysis of bacterial protein secretion system requirement for IG and JA biosynthesis regulation. Transcript accumulation of two representative IG biosynthesis genes *CYP79B2* (A) and *SOT16* (B) and of two representative JA biosynthesis genes *LOX2* (C) and *OPR3* (D) was analysed by qRT-PCR. Inoculations were performed with buffer inoculation, *D. dadantii* wild type strain (3937) and mutants altered in bacterial protein secretion systems inoculation (*prtE* type I, *outC* type II and *hrcC* type III) on Col-0 wild type *Arabidopsis.* Fold changes were normalised to 0 hpi for each inoculation condition (set to one). Error bars show the experimental values of the two biological repeats.

Figure 5A and B clearly show that IG-related gene transcript accumulation observed 30 hpi with the wild type bacterial strain was considerably lower after infection with the mutant strain with an altered protein secretion system I (T1SS prtE) and was similar to the buffer-inoculated control with the mutant strain with an altered protein secretion system II (T2SS outC). Thus, T1SS and T2SS play a role in the induction of the IG biosynthesis pathway. Conversely, inactivation of the secretion system III (T3SS hrp secretion system) leads to an approximately two-fold IG-related transcript accumulation increase as compared to the wild type strain.

Analysis of the involvement of bacterial secretion systems in the induction of biosynthesis JA genes (Figs 5C and D) leads exactly to the same conclusion: T1SS and T2SS are required to fully induce the expression of *LOX2* and *OPR3* biosynthesis genes and inactivation of T3SS slightly increases the expression levels.

The T2SS is the major secretion system both in terms of pathogenicity impact and number of secreted factors, as it is responsible, among others, for the secretion of 10 pectate lyases and 1 cellulase [47]. To clarify the dependence of IG- and JA-related gene expression on T2SS secreted factors, we analysed the expression of 2 IG- and 2 JA-related genes after Arabidopsis inoculation with the Δ*pel* mutant that is impaired in the secretion of the five major pectate lyases (PelABCDE). Figure 6 demonstrates unequivocally that these major pectate lyases are necessary for the observed inductions.

**Fig 6.**
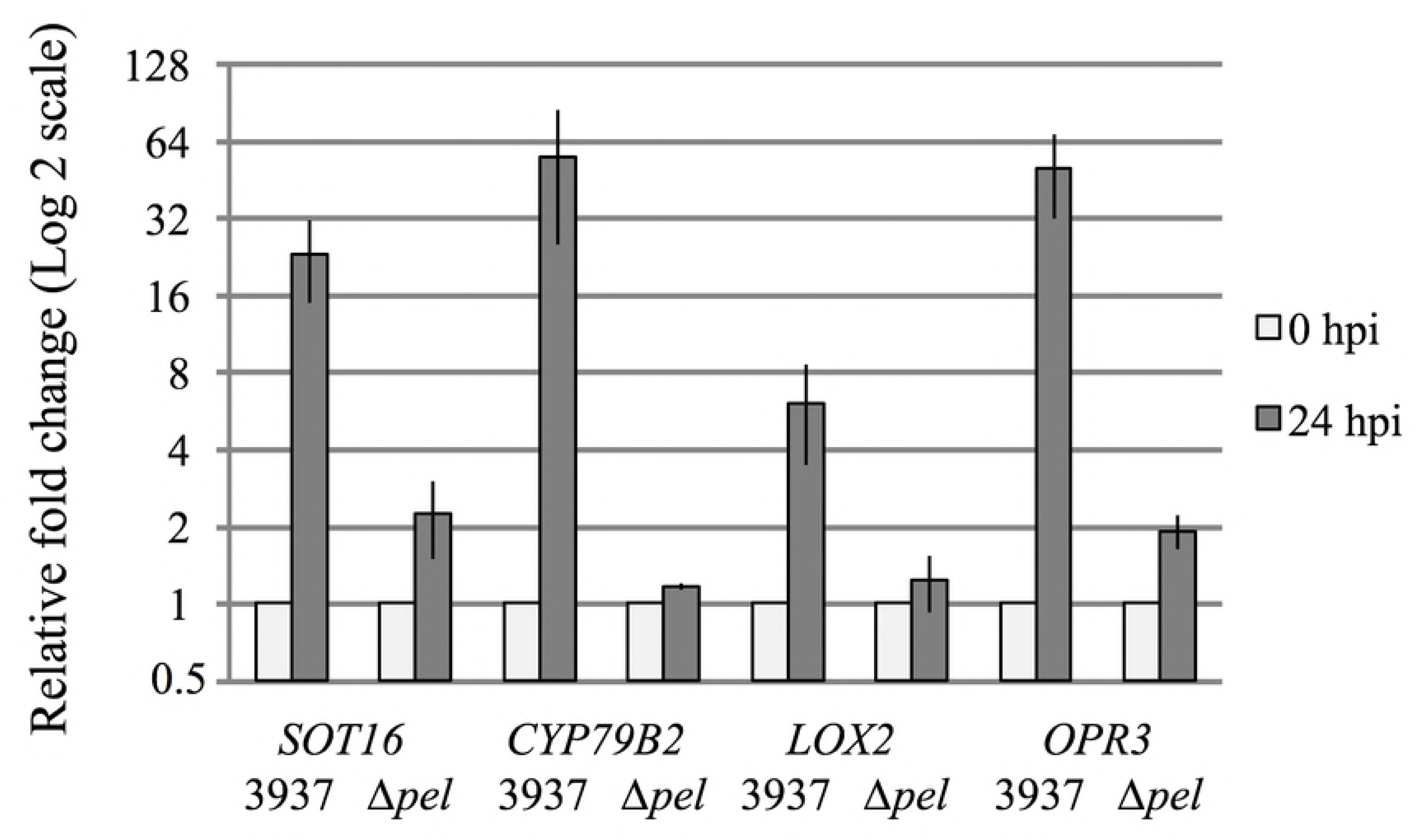
Effect of a Δpel mutation altering the five major pectate lyases synthesis on IG and JA biosynthesis induction. Expression of *CYP79B2* and *SOT16* IG biosynthesis genes, and *LOX2* and *OPR3* JA biosynthesis genes was analysed by qRT-PCR after infection by *D. dadantii* (3937) wild type strain and the Δ*pel* mutant impaired in the synthesis of the five major pectate lyases PelABCDE. Fold changes were normalised to 0 hpi values for each (set to one). Error bars show the experimental values of the two biological repeats.

### Involvement of JA and IG pathways in disease progression

In order to analyse the importance of the IG response in disease progression, we performed a kinetics analysis of symptom progression in both JA and IG-deficient *Arabidopsis* mutants as compared to the wild type parent.

The JA biosynthesis *jar1 Arabidopsis* mutant has already been briefly described as more susceptible to *D. dadantii* infection [25]. We extended this study by analysing the phenotype of two JA mutants: the JA insensitive *coi1* mutant and the biosynthetic *jar1* mutant. Plants were infected with *D. dadantii* wild-type strain, and disease progression was scored during five days using a severity symptoms scale from 0 to 3 (see legend of Fig 7) as previously described by Lebeau *et al.* [17]. In all JA mutants, we observed a significantly increased number of plants with severe symptoms (scored 3) as compared to wild-type infected plants three, four and five days post-inoculation (Fig 7A). Moreover, symptoms appear earlier (2 and 3 dpi) in the *coi1* and *jar1* mutants. This confirms that the JA pathway is involved in partial resistance of *Arabidopsis* to *D. dadantii*.

**Fig 7.**
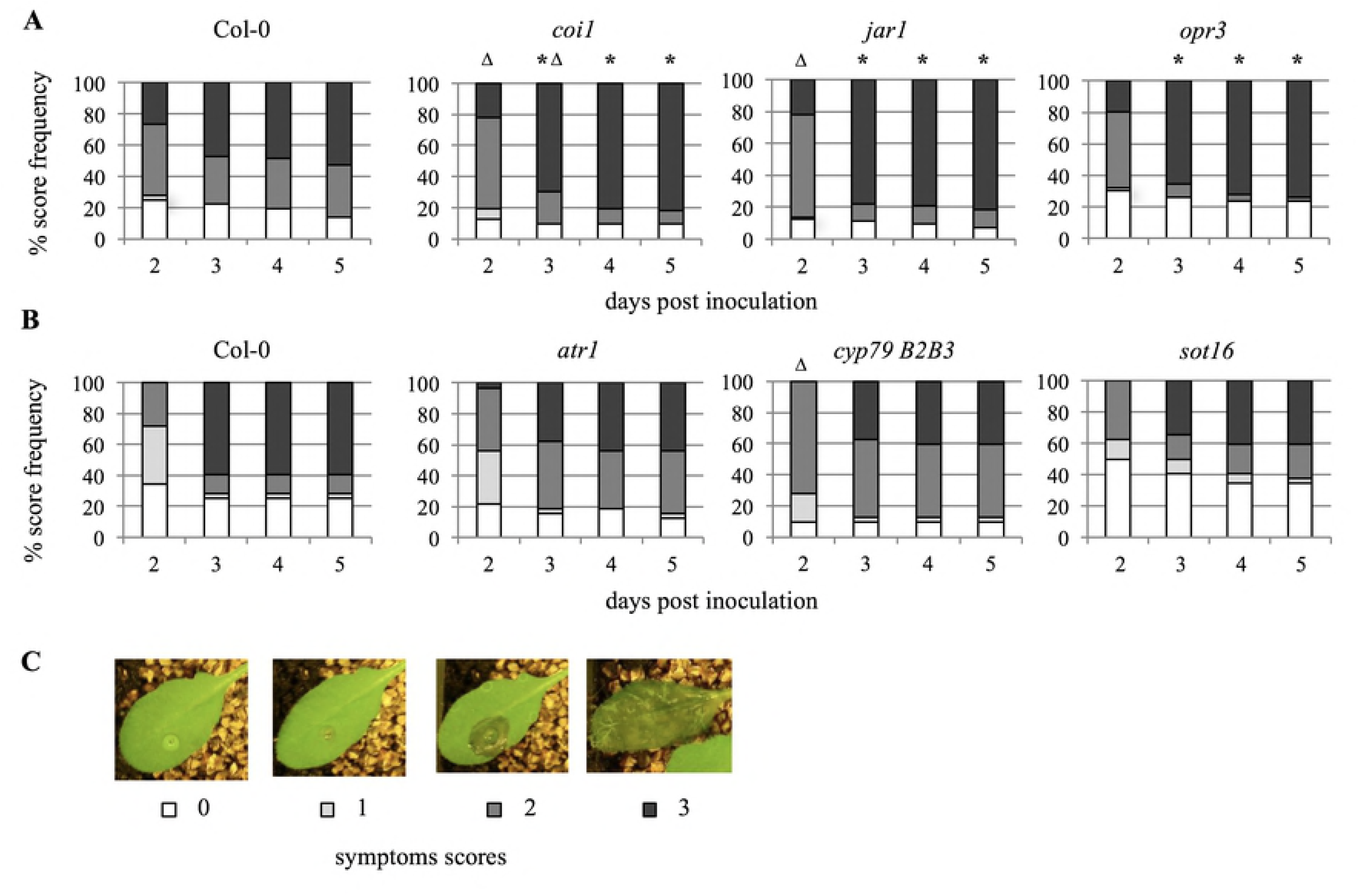
Disease severity on *Arabidopsis* wild type strain and mutants impaired in JA sensitivity or biosynthesis (A) and IG biosynthesis (B). Symptoms were scored during five days post inoculation on Col-0 and mutants using the following symptom severity scale: 0 = no symptom, 1 = maceration restricted to the site of inoculation, 2 = maceration extending from the site of inoculation, 3 = maceration covering the whole leaf (C). * and Δ indicate that score frequency of the highest degree of symptom (scored 3) and of plant with no symptom (scored 0) respectively are statistically different from the wild type one (Fisher’s exact test, two sided p-value < 0.05). Phenotypic analysis was performed on 36 individual plants and was repeated at least twice with similar results.

As IG-related genes were shown to be up-regulated during infection, we also investigated the involvement of the IG pathway in *Arabidopsis* susceptibility. Response to *D. dadantii* infection was followed in the *sot16* simple mutant and the *cyp79B2cyp79B3* double mutant, both impaired in IG biosynthesis [48, 49]. The MYB transcription factor *atr1* mutant that shows reduced IG levels [50] was also included in this study. Even if the number of plants with severe symptoms (scored 3) appeared to be reduced in IG mutants compared to the wild type, no statistically significant difference in the number of plants with these severe symptoms was observed (Fig 7B). Similarly, symptoms appearance was not statistically different, except for the *cyp79B2cyp79B3* mutant 2 dpi. Levels of intermediate symptom degree (scored 2) that are higher in the mutants compared to the wild type reflect the non-significant decreases in both symptom appearance and severity (scores 0 and 3). This led us to conclude that, even if the IG pathway is clearly activated in response to *D. dadantii* infection, this defence line has no significant impact on disease progression.

## Discussion

To study the early plant response to *D. dadantii* infection, we performed an *Arabidopsis* whole-genome transcriptome analysis during the early phase of infection. We used an infection procedure - a rapid immersion in bacterial suspension avoiding plant wounding - much closer to a natural process than an infiltration with a high-density bacterial suspension. This strategy was chosen to prevent a massive transcriptome reprogramming due to rapidly spreading maceration, with considerable cell death and dramatic perturbation of plant homeostasis. As expected, the expression of only a limited number of genes was modulated during the early stages of infection and several of them were strongly associated with plant defence and interaction with the environment. To our knowledge, this study is the first *Arabidopsis* transcriptome analysis at a whole-genome level in response to soft rot enterobacterales infection. It substantially diverges from other transcriptome studies involving for example the T3SS-dependent bacterial *P. syringae* or the necrotrophic fungus *B. cynerea*, for which one or two thousands of genes are modulated [51, 52]. It enabled us to show a coordinated transcriptional induction of genes involved in the metabolism of tryptophan and the related indole glucosinolate secondary metabolites (Fig 2). Moreover, our study also revealed the modulation of JA-related genes expression.

The coordinated activation of IG and JA pathways after *D. dadantii* infection evokes a role of jasmonates in *D. dadantii*-provoked IG response. Indeed, previous studies reported that IG production and IG-related genes expression were mediated, at least partially, by jasmonates, even if the experimental contexts (bacterial elicitors or MeJA treatments, chewing insects attack) were different from ours [53–59]. Our results however depict a more subtle situation. First, induction of the *CYP79B2* gene encoding (with *CYP79B3*) the bottleneck enzyme of tryptophan-derived secondary metabolism [60] was shown to be independent of the JA pathway while the induction of the downstream *SOT16* biosynthesis gene was dependent on the JA receptor COI1 (Fig 4). The occurrence of differential regulations along a biosynthetic pathway is not unusual and could be related to the involvement of this pathway in a wide metabolic network. Indeed, indole acetaldoxime - the product of CYP79B2 activity - serves as a branching point for indole glucosinolates, IAA and camalexin biosynthesis (for review see [61]). From that perspective, it can be easily conceived that a regulation of genes involved in a final synthesis step would be more efficient to control total IG contents without disturbing others pathways. Second, *SOT16* gene induction was shown to be dependent on the JA receptor COI1, but not on the key JA biosynthesis jasmonate-amido synthetase enzyme, encoded by the *JAR1* gene. Such contrasted effects of mutations inactivating *COI1* and JA biosynthesis genes have often been described in other systems. For example, the *jar1* mutant was shown to have little or no effect on wound induction of JA-regulated genes such as *PDF1.2*, although JA–Ile is known to be a key signal for COI1 activity [62]. Furthermore, stress-related response genes that specifically respond to the JA-precursor oxo-phytodienoate OPDA but not to JA have been identified [63, 64]. More recently, phytoprostane oxylipins - prostaglandin/jasmonate-like products of nonenzymatic lipid peroxidation - were shown to regulate specific plant responses in a COI1-dependent manner but independently of jasmonate biosynthesis [65]. This multiplicity of bioactive JA compounds and their different modes of action could allow plants to respond specifically and flexibly to pathogen attack [66].

Brader *et al.* [54] reported the activation of Trp and IG biosynthetic pathways in *Arabidopsis* by culture filtrates of *Pectobacterium*. This prompted us to analyse JA and IG-related gene activation after infection with *D. dadantii* mutants impaired in the major protein secretion systems involved in virulence to find out if a secreted virulence factor could trigger this gene induction. We showed that the T1SS (PrtE) and T2SS (OutC) were individually necessary to fully induce both pathways (Fig 5). Notably, induction impairment was more severe after infection by the *outC* mutant than by the *prtE* mutant for both pathways. Of the hundred proteins secreted by *D. dadantii*, pectinases are the predominant virulence factors secreted by the T2SS, as revealed by the phenotype of the *outC* mutant that is totally unable to provoke maceration symptoms [9]. We therefore extended our analysis to the Δ*pel* mutant that is impaired in the synthesis of the five major pectate lyases PelABCDE. This analysis showed that this limited set of five pectate lyases was required to trigger the observed inductions of both JA and IG pathways (Fig 6).

These inductions might be caused either by a direct effect of proteins or by the enzymatic release of signalling oligogalacturonides. It indicates furthermore that *D. dadantii* does not exhibit MAMPs that can induce these pathways. Such MAMP-independent induction has already been reported for example in the *Erwinia amylovora/Malus spp.* pathosystem [67]. Interestingly, this MAMP-independent induction contrasts with the induction of all known IG biosynthetic genes via the MYB51 transcription factor in *Arabidopsis* triggered by the *P. syringae-*derived Flg22 flagellin peptide fragment [68]. To that respect, it should be noted that the *D. dadantii* flagellin FliC shares only 63.3% identity in a 14 aa overlap with the Flg22 peptide and amino acid substitutions concern some amino acids which are known to be important for elicitor activity [69]. This could explain the lack of induction of the IG pathway by *D. dadantii* flagellin. We can envision that the lack of some plant responses to *D. dadantii* MAMPs would represent an adaptive advantage for the pathogen during the early asymptomatic stage, the subsequent activation of plant responses during the following symptomatic stage possibly occurring too late to be efficient.

Compared to the much weaker induction observed with T1SS and T2SS mutants, induction of both JA- and IG-related genes after infection by the T3SS *hrcC* mutant was about 2 times higher than after infection by the wild type 30 hpi (Fig 5). It is well documented that T3SS of plant pathogenic bacteria injects effector proteins that suppress basal defence (for review, see [70]) and, in different pathosystems, T3SS-defective bacteria were found to highly activate basal defences (for review, see [71, 72]). Furthermore, a strong T3SS-dependent down-regulation of the JA pathway was observed in leaves of the susceptible *Malus spp*. genotype following *E. amylovora* infection [67]. Although T3SS only plays a marginal role in *D. dadantii* virulence - this system being important only at low inoculum or on semi-tolerant plants [12, 16] - the stronger induction of JA and IGs-related genes observed with the *hrcC* mutant could reveal an involvement of type III effectors in the down-regulation of JA-related basal defences.

IGs participate, with aromatic and aliphatic glucosinolates, in the capparales-specific plant defence glucosinolate-myrosinase system (for review, see [73]) that is often referred as the mustard oil bomb. This system, effective against most herbivorous insects and against necrotrophic fungi, is based on the compartmentalization of glucosinolates and matching hydrolases, called myrosinases. Glucosinolates form a major group of secondary metabolites, based on a core structure of a ß-D-thioglucose group linked to a sulfonated aldoxime moiety. When plant tissue is damaged, glucosinolates come into contact with plant myrosinases. Myrosinases remove the ß-glucose moiety from glucosinolates, leading to the formation of an unstable intermediate, and, finally, to a variety of toxic breakdown products (Fig 2) involved in resistance to pathogens including microorganisms, nematodes and insects (for review, see [74]). More recently, the role of IG breakdown products was extended to plant defence signalling [68].

The specific induction of IGs biosynthesis and degradation genes taking place after inoculation of *Arabidopsis* by *D. dadantii* evokes a role of IGs in the *Arabidopsis* defence response. While glucosinolates have been shown to play a clear role in defence against herbivory and fungal pathogens [73, 75], such evidence is poorly documented concerning the role of IGs in plant-bacteria interactions. The only reports on an antibacterial effect of IGs - including on *D. dadantii* - are *in vitro* studies using IGs breakdown products [54, 76, 77]. However, these studies minimize the complexity of the glucosinolate response, as illustrated by the diversity of glucosinolate species (more than 40 molecules, of which 4 known IGs) and of nitrile specifier enzymes that generate various nitrile species in conjunction with myrosinases. In addition, *in vitro* studies do not take into account well known bacterial adaptations to various plant generated stresses encountered during infection, such as acidic and oxidative stresses. Thus, we could potentially imagine a bacterial adaptive response to toxic IG-breakdown products.

The phenotypic analysis of three Arabidopsis mutants impaired in IGs biosynthesis (*atr1*, *cyp79B2cyp79B3* and *sot16*) did not reveal any significant difference in symptom severity after *D. dadantii* inoculation as compared to the wild type (Fig 7B). This indicates that IGs do not play *in planta* a major role in plant defence against *D. dadantii*. In contrast, the involvement of the JA pathway in limiting symptom progression during infection previously reported by Fagard *et al.* [25] was clearly confirmed by the phenotypes of the three JA mutants we analysed (Fig 7A). While IG defence response has been demonstrated to be effective during chewing insects attack or necrotrophic fungi infection, previous studies on model phytopathogenic bacteria (*P. syringae, E. carotovora, Pectobacterium*…) do not report any clear evidence of an effect of IG production on disease progression. Our results illustrate what may be a general trait of bacterial infections: a clear and unambiguous induction of the IG pathway, balanced by the lack of effectiveness of this defence response against bacterial pathogens. Such an inefficient massive defence response to *D. dadantii* was already observed by Fagard *et al*. [25], who described the induction of the SA pathway and an absence of sensitivity phenotype of the corresponding mutants.

In conclusion, our transcriptomic analysis of *Arabidopsis* responses to *D. dadantii* infection revealed the activation of two pathways related to plant defence, the IG and JA pathways. The activation of some of IG biosynthesis genes requires a functional COI1, but is independent of jasmonate and its derivative conjugate JA-Ile synthesis. Mutant analysis revealed however that, while JA signalling is clearly involved in defence against this pathogen, IGs production does not appear to hinder *D. dadantii* virulence. The impaired activation of these pathways by T1SS and T2SS bacterial mutants highlights that *D. dadantii* MAMPs recognition has no effect on this induction. This could explain the weakness of *Arabidopsis* response to *D. dadantii* infection, in term of number of modulated genes, as compared to infection by other bacteria, necrotophic fungi or chewing insects. This camouflage strategy during the early stage of infection could be one element explaining the adaptive success of this bacterial pathogen, as exemplified by the host broadness and the lack of specific and highly efficient defence.

## Acknowledgments

Transcriptome profiling using CATMA arrays was supported by a grant from the French National Institute for Agricultural Research (INRA).

We thank Jean-Pierre Renou for CATMA array experiment design and data analysis, Caroline Kunz and Marie-Anne Barny for critical reading of the manuscript and Emma Rochelle-Newall for English corrections. We thank J. Bender for *cyp79B2cyp79B3* and *atr1* mutants, M. Fagard for *coi1* mutants.

## Data deposition

Microarray data were deposited at Gene Expression Omnibus database (accession number GSE11793) and at CATdb (project RA06-04_BOS1) according to the “Minimum Information About a Microarray Experiment” standards.

## Supporting information

**Table S1. Bacterial strains, plant lines and primers used in this study.** Primer efficiency (E) for qPCR was calculated from standard curve slope according to E = (10 ^(−1/slope)^ - 1) x 100. Primers were selected in a range of 100±5 % efficiency.

**Table S2. List of total *Arabidopsis* genes modulated during *D. dadantii* infection 12 or 24 hpi.** Fold change ratios (infected plants *vs* buffer control) are expressed in log2 scale. When p-value were higher than 0.05, fold change ratios were shaded in light grey. Red scale bar: up-regulated genes; green scale bar: down-regulated genes. The threshold for modulated genes was fixed to ± 0.5 (log2 scale).

## Abbreviations

IGs: indole glucosinolates
ROS: reactive oxygen species
PR: pathogenesis-related
ET: ethylene
T#SS: type # secretion system
PCD: plant cell death
hpi: hours post inoculation
MAMPs: microbe-associated molecular patterns

